# Predictive Neural Computations Support Spoken Word Recognition: Evidence from MEG and Competitor Priming

**DOI:** 10.1101/2020.07.01.182717

**Authors:** Yingcan Carol Wang, Ediz Sohoglu, Rebecca A. Gilbert, Richard N. Henson, Matthew H. Davis

## Abstract

Human listeners achieve quick and effortless speech comprehension through computations of conditional probability using Bayes rule. However, the neural implementation of Bayesian perceptual inference remains unclear. Competitive-selection accounts (e.g. TRACE) propose that word recognition is achieved through direct inhibitory connections between units representing candidate words that share segments (e.g. *hygiene* and *hijack* share /haidʒ/). Manipulations that increase lexical uncertainty should increase neural responses associated with word recognition when words cannot be uniquely identified. In contrast, predictive-selection accounts (e.g. Predictive-Coding) proposes that spoken word recognition involves comparing heard and predicted speech sounds and using prediction error to update lexical representations. Increased lexical uncertainty in words like *hygiene* and *hijack* will increase prediction error and hence neural activity only at later time points when different segments are predicted. We collected MEG data from male and female listeners to test these two Bayesian mechanisms and used a competitor priming manipulation to change the prior probability of specific words. Lexical decision responses showed delayed recognition of target words (*hygiene*) following presentation of a neighbouring prime word (*hijack*) several minutes earlier. However, this effect was not observed with pseudoword primes (*higent*) or targets (*hijure*). Crucially, MEG responses in the STG showed greater neural responses for word-primed words *after* the point at which they were uniquely identified (after /haidʒ/ in *hygiene*) but not *before* while similar changes were again absent for pseudowords. These findings are consistent with accounts of spoken word recognition in which neural computations of prediction error play a central role.

**Significance Statement:** Effective speech perception is critical to daily life and involves computations that combine speech signals with prior knowledge of spoken words; that is, Bayesian perceptual inference. This study specifies the neural mechanisms that support spoken word recognition by testing two distinct implementations of Bayes perceptual inference. Most established theories propose direct competition between lexical units such that inhibition of irrelevant candidates leads to selection of critical words. Our results instead support predictive-selection theories (e.g. Predictive-Coding): by comparing heard and predicted speech sounds, neural computations of prediction error can help listeners continuously update lexical probabilities, allowing for more rapid word identification.

## Introduction

In daily conversation, listeners identify ~200 words/minute (Tauroza & Allison, 1990) from a vocabulary of ~40,000 words (Brysbaert et al., 2016). This means that they must recognise 3-4 words/second and constantly select from sets of transiently ambiguous words (e.g. *hijack and hygiene* both begin with /haidʒ/). Although it is recognised that humans achieve word recognition by combining current speech input with its prior probability using Bayes theorem (Norris & McQueen, 2008; Davis & Scharenborg, 2016; Gwilliams & Davis, in press), the underlying neural implementation of Bayesian perceptual inference remains unclear (Aitchison & Lengeyl, 2017).

Here, we test two computational accounts of spoken word recognition that both implement Bayes rules. In competitive-selection accounts (e.g. TRACE, McClelland & Elman, 1986, Figure 1A), word recognition is achieved through within-layer lateral inhibition between neural units representing similar words. By this view, *hijack* and *hygiene* compete for identification such that an increase in probability for one word inhibits units representing other similar-sounding words. Conversely, predictive-selection accounts (e.g. Predictive-Coding, Davis & Sohoglu, 2020) suggest that word recognition is achieved through computations of prediction error (Figure 1D). On hearing transiently ambiguous speech like /haidʒ/, higher-level units representing matching words make contrasting predictions (/æk/ for *hijack*, /i:n/ for *hygiene*).Prediction error (the difference between sounds predicted and actually heard) provides a signal to update word probabilities such that the correct word can be selected.

**Figure 1.**
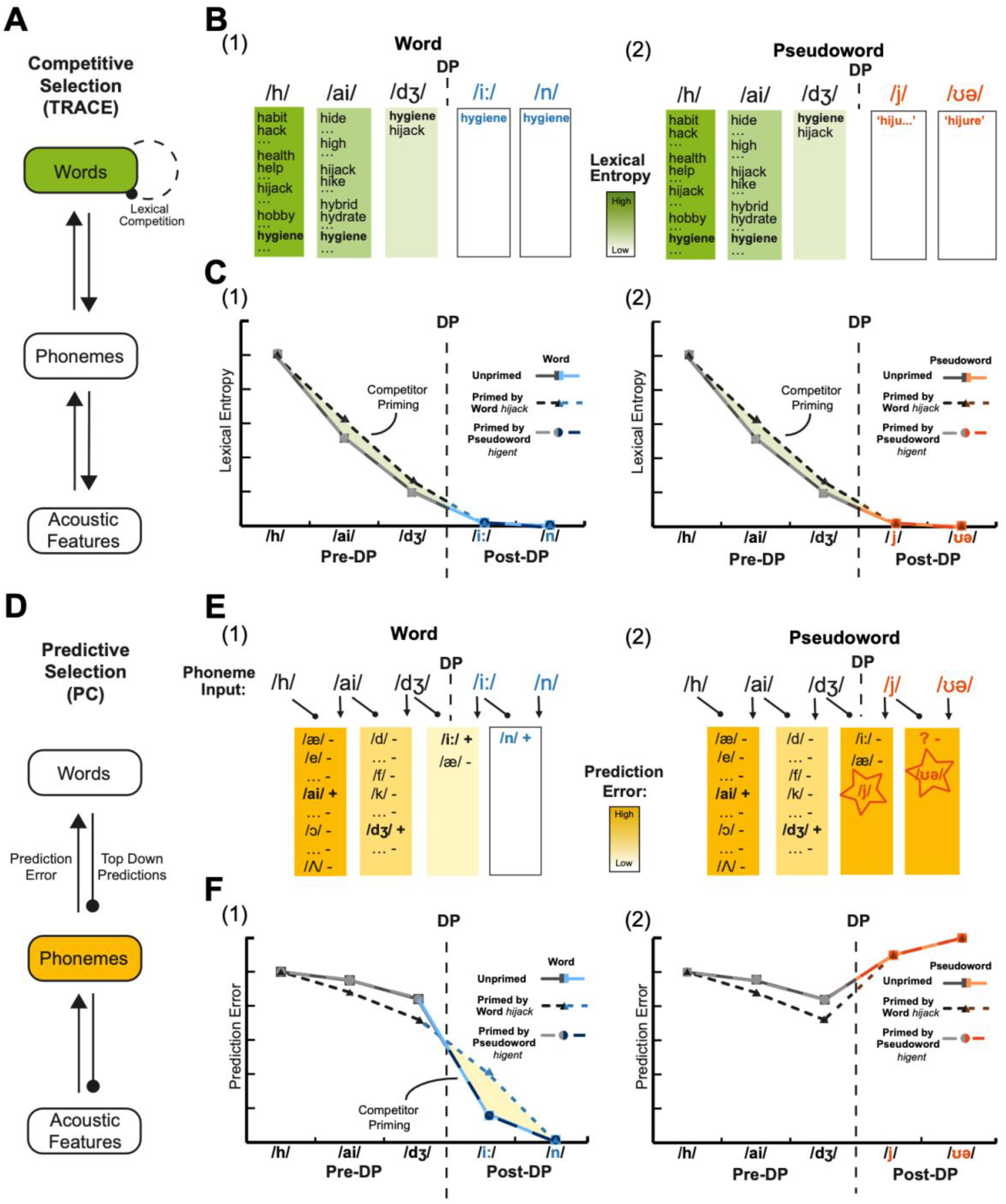
Illustration of neural predictions based on competitive-selection and predictive-selection models respectively for recognition of a word (*hygiene*) or pseudoword (*hijure*) that is unprimed or primed by a similar-sounding word (*hijack*) or pseudoword (*higent*). ***A.*** In a competitive-selection model, such as TRACE (McClelland & Elman, 1986), word recognition is achieved through within-layer lexical competition. ***B.*** Illustration of the competitive-selection procedure for word (e.g. *hygiene*) and pseudoword (e.g. *hijure*) recognition. Phoneme input triggers the activation of multiple words beginning with the same segments, which compete with each other until one word is selected. No word can be selected when hearing a pseudoword, though it would be expected that lexical probability (although not lexical entropy) should be greater for words than for pseudowords. ***C.*** Illustration of neural predictions based on lexical entropy. Lexical entropy gradually reduces to zero as more speech is heard. Before the deviation point (hereafter DP) at which the prime (*hijack*) and target (*hygiene*) diverge, these items are indistinguishable, and competitor priming should transiently increase lexical entropy (shaded area). After the DP, competitor priming should not affect entropy since prime and target words can be distinguished. ***D.*** In a predictive-selection model such as the Predictive-Coding account (PC, Davis & Sohoglu, 2020), words are recognised by minimising prediction error, which is calculated by subtracting the predicted segments from the current sensory input. ***E.*** Illustration of the predictive-selection procedure during word (e.g. *hygiene*) and pseudoword (e.g. *hijure*) recognition. Speech input evokes predictions for the next segment (based on word knowledge as in B), which is then subtracted from the speech input and used to generate prediction errors that update lexical predictions (+ shows confirmed predictions that increase lexical probability, - shows disconfirmed predictions that decrease lexical probability). ***F.*** Illustration of neural predictions based on segment prediction error. Before the DP, priming of initial word segments should strengthen predictions and reduce prediction error. There will also be greater mismatch between predictions and heard speech for competitor-primed words and hence primed words should evoke greater prediction error than unprimed words (shaded area). This increased prediction error should still be less than that observed for pseudowords, which should evoke maximal prediction error regardless of competitor priming due to their post-DP segments being entirely unpredictable.

In this study, we used the competitor priming effect (Monsell & Hirsh, 1998; Marsolek, 2008), which is directly explicable in Bayesian terms, i.e. the recognition of a word (*hygiene*) is delayed if the prior probability of a competitor word (*hijack*) has been increased due to an earlier exposure. This delay could be due to increased lateral inhibition (competitive-selection) or greater prediction error (predictive-selection). Thus, similar behavioural effects of competitor priming are predicted by two distinct neural computations (Spratling, 2008). To distinguish them, it is critical to investigate neural data that reveals the direction, timing and level of processing at which competitor priming modulates neural responses. Existing neural data remains equivocal with some evidence consistent with competitive-selection (Bozic et al., 2010; Okada & Hickok, 2006), predictive-selection (Gagnepain et al, 2012), or both mechanisms (Brodbeck et al., 2018; Donhauser et al., 2019). We followed these studies in correlating two computational measures with neural activity: lexical entropy (competitive-selection) and segment prediction error (or phoneme surprisal, for predictive-selection).

Here, we used MEG to record the location and timing of neural responses during spoken words recognition in a competitor priming experiment. Pseudowords (e.g. *hijure*) were included in our analysis to serve as a negative control for competitor priming, since existing research found that pseudowords neither produce nor show this effect (Monsell & Hirsh, 1998). We compared items with the same initial segments (words *hygiene, hijack*, pseudowords *hijure, higent* share /haidʒ/) and measured neural and behavioural effects concurrently to link these two effects for single trials.

While lexical entropy and prediction error are correlated for natural speech, this competitor priming manipulation allows us to make differential predictions as illustrated in Figure 1. Specifically: (1) before the deviation point (DP, the point at which similar-sounding words diverge), competitor priming increases lexical entropy and hence neural responses (Figure 1B&C Pre-DP). Such responses might be observed in inferior frontal regions (Zhuang et al., 2011) and posterior temporal regions (Prabhakaran et al., 2006). However, prediction error will be reduced for pre-DP segments, since heard segments are shared and hence more strongly predicted (Figure 1E&F Pre-DP). This should be reflected in the superior temporal gyrus (STG, Sohoglu & Davis, 2016). (2) After the DP, predictive-selection but not competitive-selection accounts propose that pseudowords evoke greater signals in the left-STG, since they evoke maximal prediction errors (Figure 1E&F Pseudoword, Post-DP). (3) Furthermore, in predictive-selection theories, competitor priming is associated with an increased STG response to post-DP segments due to enhanced prediction error caused by mismatch between primed words (predictions) and heard speech (Figure 1E&F Word, Post-DP).

## Materials and Methods

### Participants

Twenty-four (17 female, 7 male) right-handed, native English speakers were tested after giving informed consent under a process approved by the Cambridge Psychology Research Ethics Committee. This sample size was selected based on previous studies measuring similar neural effects with the same MEG system (Gagnepain et al. 2012; Sohoglu & Davis, 2016; Sohoglu et al. 2012, etc.). All participants were aged 18-40 years and had no history of neurological disorder or hearing impairment based on self-report. Two participants’ MEG data were excluded from subsequent analyses respectively due to technical problems and excessive head movement, resulting in 22 participants in total. All recruited participants received monetary compensation.

### Experimental Design

To distinguish competitive- and predictive-selection accounts, we manipulated participants’ word recognition process by presenting partially mismatched auditory stimuli prior to targets. Behavioural responses and MEG signals were acquired simultaneously. Prime and target stimuli pairs form a repeated measures design with two factors (lexicality and prime type). The lexicality factor has 2 levels: word and pseudoword, while the prime type factor contains 3 levels: unprimed, primed by same lexical status, primed by different lexical status. Hence the study is a factorial 2 x 3 design with 6 conditions: unprimed word (*hijack*), word-primed word (*hijack-hygiene*), pseudoword-primed word (*basef-basis*), unprimed pseudoword (*letto*), pseudoword-primed pseudoword (*letto-lettan*), word-primed pseudoword (*boycott-boymid*). Prime-target pairs were formed only by stimuli sharing the same initial segments. Items in the two unprimed conditions served as prime items in other conditions and they were compared with target items (Figure 2A).

**Figure 2.**
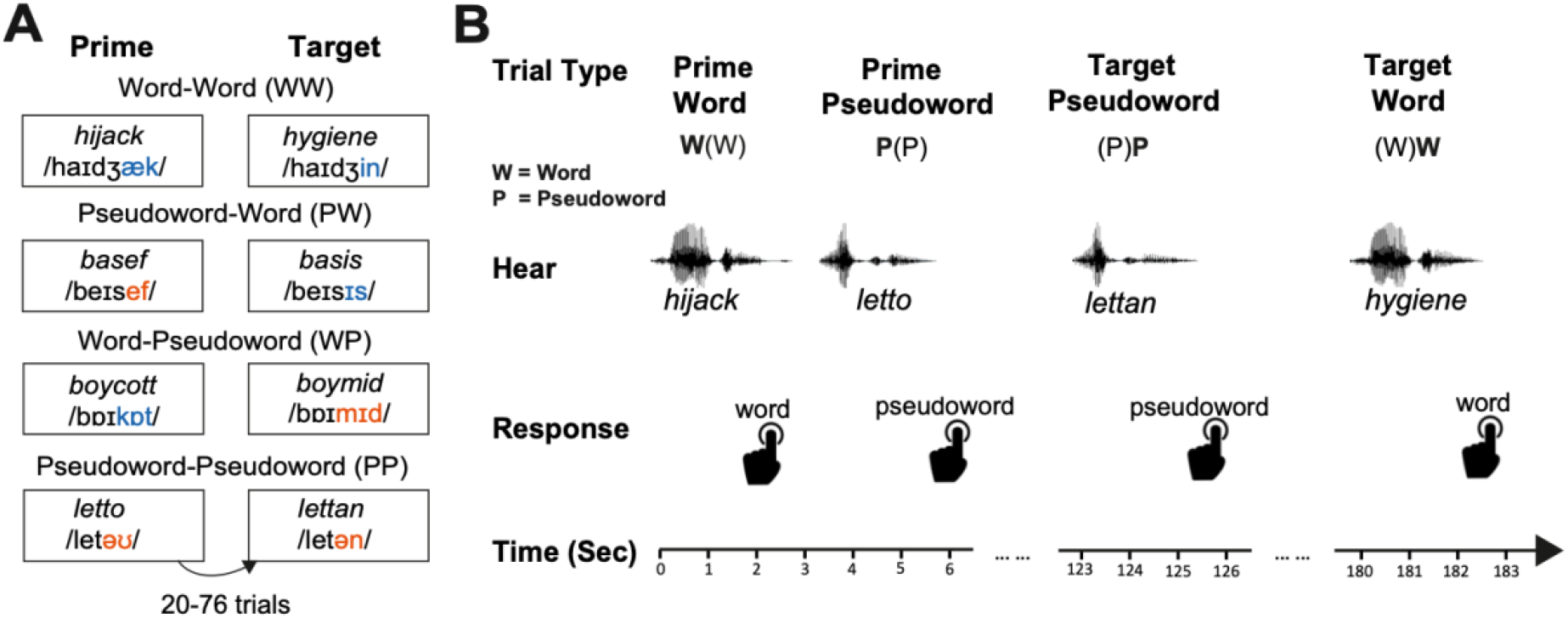
Experimental design. ***A.*** Four different types of prime-target pairs. Each pair was formed by two stimuli from the same quadruplet, separated by between 20 to 76 trials of items that do not share the same initial segments. ***B.*** Lexical decision task. Participants made lexicality judgments to each item they heard via left hand button-press. The response time was recorded from the onset of the stimuli. As shown, items within each quadruplet are repeated after a delay of ~1-4 minutes following a number of other intervening stimuli.

The experiment used a lexical decision task (Figure 2B) implemented in MATLAB through Psychtoolbox-3 (Kleiner et al. 2007), during which participants heard a series of words and pseudowords while making lexicality judgments to each stimulus by pressing buttons using their left index and middle fingers only, with the index finger pressing one button indicating word and the middle finger pressing the other button indicating pseudoword. 344 trials of unique spoken items were presented every ~3 seconds in two blocks of 172 trials, each block lasting approximately 9 minutes. Each prime-target pair was separated by 20 to 76 trials of items that do not start with the same speech sounds, resulting in a relatively long delay of ~1-4 minutes between presentations of phonologically-related items. This delay was chosen based on Monsell and Hirsh (1998), who suggest that it prevents strategic priming effects (Norris et al. 2002). Stimuli from each of the quadruplets were Latin-square counterbalanced across participants, i.e. stimulus quadruplets that appeared in one condition for one participant were allocated to another condition for another participant. The stimulus sequences were pseudo-randomised using Mix software (van Casteren & Davis, 2006), so that the same type of lexical status (word/pseudoword) did not appear successively on more than 4 trials.

### Stimuli

The stimuli consisted of 160 sets of four English words and pseudowords, with durations ranging from 372 to 991 ms (M = 643, SD = 106). Each set contained 2 words (e.g. *letter, lettuce*) and 2 phonotactically-legal pseudowords (e.g. *letto, lettan*)that share the same initial segments (e.g. /let/) but diverge immediately afterwards.

We used polysyllabic word pairs (M_syllable_ = 2.16, SD_syllable_ =0.36) instead of monosyllabic ones in our experiments so as to identify a set of optimal lexical competitors that are similar to their prime yet dissimilar from all other items. All words were selected from the CELEX database (Baayen et al., 1993). Their frequencies were taken from SUBTLEX UK corpus (Van Heuven et al., 2014) and restricted to items under 5.5 based on log frequency per million word (Zipf scale, Van Heuven et al., 2014). In order to ensure that any priming effect was caused purely by phonological but not semantic similarity, we also checked that all prime and target word pairs have a semantic distance of above 0.7 on a scale from 0 to 1 based on the Snaut database of semantic similarity scores (Mandera et al., 2017), such that morphological relatives (e.g. darkly/darkness) were excluded.

All spoken stimuli were recorded onto a Marantz PMD670 digital recorder by a male native speaker of southern British English in a sound-isolated booth at a sampling rate of 44.1 kHz. Special care was taken to ensure that shared segments of stimuli were pronounced identically (any residual acoustic differences were subsequently eliminated using audio morphing as described below).

The point when items within each quadruplet begin to acoustically differ from each other is the deviation point (hereafter DP, see Figure 3A). Pre-DP length ranged from 150 to 672 ms (M = 353, SD = 96), while post-DP length ranged from 42 to 626 ms (M = 290, SD = 111, see Figure 3B). Epochs of MEG data were time-locked to the DP. Using phonetic transcriptions (phonDISC) in CELEX, the location of the DP was decided based on the phoneme segment at which items within each quadruplet set diverge (M_seg_=3.53, SD_seg_=0.92). To determine when in the speech files corresponds to the onset of the first post-DP segment, we aligned phonetic transcriptions to corresponding speech files using the WebMAUS forced alignment service (Kisler et al., 2017; Schiel, 1999). In order to ensure that the pre-DP portion of the waveform was acoustically identical, we cross-spliced the pre-DP segments of the 4 stimuli within each quadruplet and conducted audio morphing to combine the syllables using STRAIGHT (Kawahara, 2006) implemented in MATLAB. This method decomposes speech signals into source information and spectral information, and permits high quality speech re-synthesis based on modified versions of these representations. This enables flexible averaging and interpolation of parameter values that can generate acoustically intermediate speech tokens (see Rogers & Davis, 2017, for example). In the present study, this method enabled us to present speech tokens with entirely ambiguous pre-DP segments, and combine these with post-DP segments without introducing audible discontinuities or other degradation in the speech tokens. This way, phonological co-articulation in natural speech was reduced to the lowest level possible at the DP, hence any cross-stimuli divergence evoked in neural responses can only be caused by post-DP deviation.

**Figure 3.**
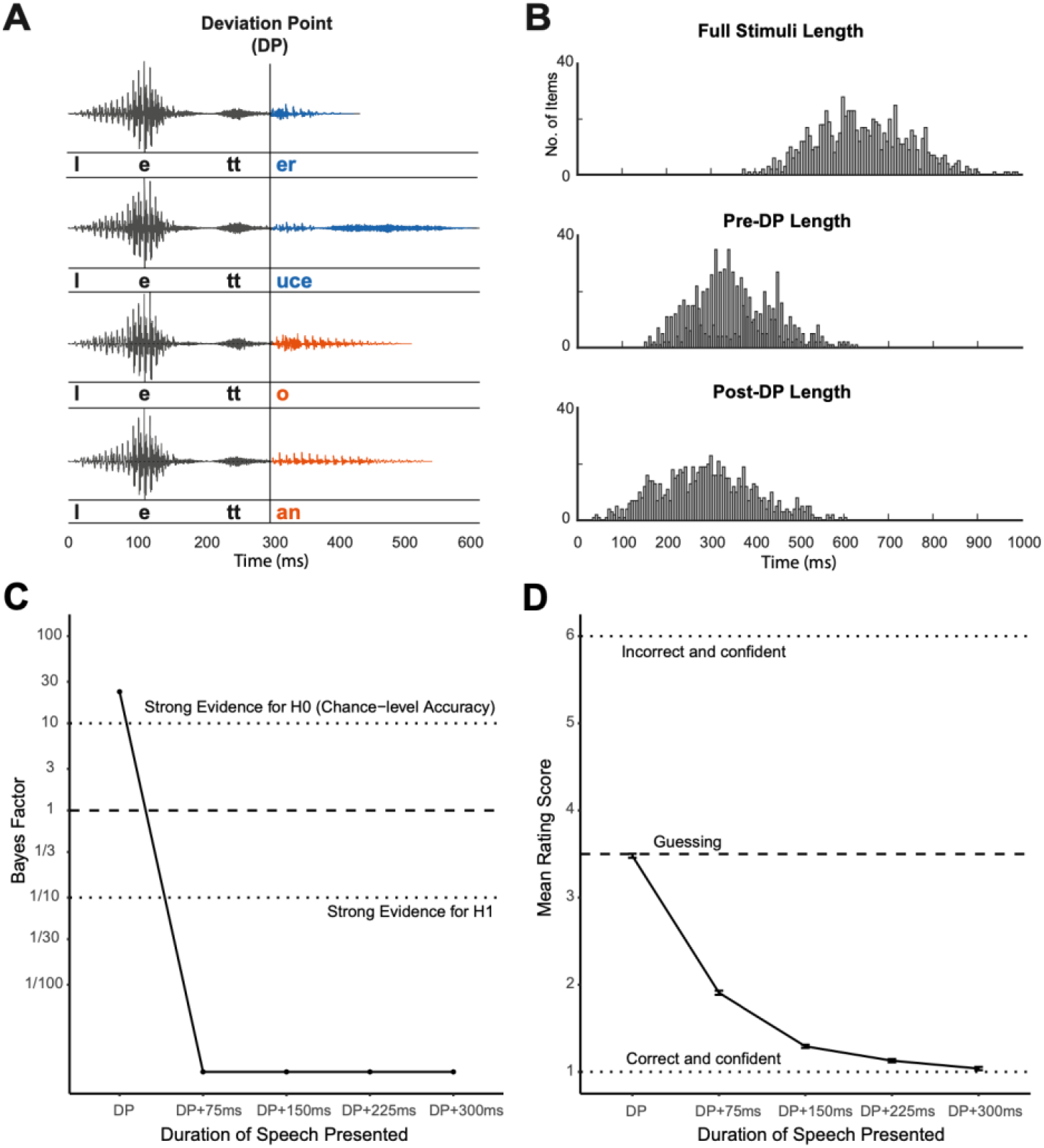
Stimuli and post-test gating study results. ***A.*** Stimuli within the same quadruplet have identical onsets in STRAIGHT parameter space (Kawahara, 2006) and thus only diverge from each other after the deviation point (DP). MEG responses were time-locked to the DP. ***B.*** Stimuli length histogram. ***C.*** Bayes factor for chance level accuracy (BF01) at each post-DP alignment point of the stimuli in the post-test gating study. ***D.*** Mean rating score at each post-DP alignment point of the stimuli in the gating study.

### Post-test Gating Study

As encouraged by a reviewer, we conducted a post-test perceptual experiment using a gating task in order to confirm that the cross-splicing and morphing of our stimuli worked as expected. This experiment used a gating task implemented in JavaScript through JSpsych (de Leeuw, 2015). During the experiment, auditory segments of all 160 pairs of words used in the MEG study were played. Twenty British English speakers were recruited through Prolific Academic online with monetary compensation. The sample size was selected based on a similar gating study conducted by Davis et al. (2002). Participants were evenly divided into two groups, one group were presented with 160 stimuli words with different pre-DP segments (e.g. *hygiene*), while the other group were presented with the other paired 160 stimuli (e.g. *hijack*). Therefore, participants only ever heard one of the two items in each pair. Stimuli segments of each word item consist of the pre-DP segment and, depending on the stimuli length, also longer segments that are 75ms, 150ms, 225ms and 300ms post DP. The segments of each word were presented in a gating manner, with the shortest segment played the first and the full item played at the end. After hearing each segment (e.g. /haidʒ/), participants were also presented with the writing of the word (e.g. *hygiene*) that contained the segment and the other paired word that shared the same pre-DP segment (e.g. *hijack*) on the screen. We asked the participants to choose which item the auditory segment matches and indicate their confidence from a rating scale of 1 to 6, with 1 representing being very confident that the item is the one on the left and 6 representing being very confident that the item is the one on the right, while 3 and 4 representing guessing the possible item. In order to avoid potential practice effect, we also added 40 filler stimuli that are identifiable on initial presentation.

Given our goal of assessing whether there is any information to distinguish the words prior to the divergence point, we needed to adopt an analysis approach that could confirm the null hypothesis that no difference exists between perception of the shared first syllable of word pairs like *hijack* and *hygiene*. We therefore analysed the results using Bayesian methods which permit this inference. Participants’ response accuracy was analysed using mixed-effect logistic regression and confidence rating scores were analysed using mixed-effect linear regression using the brms package (Bürkner, 2017) implemented in R. Response scores were computed in a way such that correct and most confident responses were scored 1, while incorrect and most confident responses were scored 6 and so on. Participants and items were included as random factors of the models and there was no fixed factor since we are only interested in the intercepts, whose estimates indicate the logit transformed proportion of correctness in the logistic model and the mean rating in the linear model respectively. We chose weakly informative priors for each model and conducted Bayes Factor analyses through the Savage-Dickey density ratio method (Wagenmakers et al., 2010). Model estimate, standard error, lower and upper boundary of 95% credible interval (CI) are also reported.

When checking our data, we found that 16 pairs of word items were not morphed correctly, hence the spectral information of the pre-DP segments of these word pairs were not exactly the same and some of them diverged acoustically before the DP due to coarticulation. Therefore, we excluded these items from analyses of the gating data and confirmed that excluding these items did not modify the interpretation or significance of the MEG or behavioral results reported in the paper.

As shown in Figure 3C, we found that when gating segments ended at the DP, Bayes factor provides strong evidence in favour of the null hypothesis, chance-level accuracy (i.e. proportion of correct responses is 0.5), β = 0.04, *SE* = 0.08, lCI = −0.11, uCI = 0.20, BF01 = 23.04. This indicates that participants could not predict the full stimuli based on hearing the pre-DP segments. On the other hand, the Bayes factor at later alignment points is close to 0, providing extremely strong evidence for the alternative hypothesis that the proportion of correct responses is higher than 0.5 (75ms post-DP: β = 3.41, *SE* = 0.22, lCI = 2.99, uCI = 3.85, BF01 < 0.01; 150ms post-DP: β = 6.26, *SE* = 0.56, lCI = 5.24, uCI = 7.41, BF01 < 0.01; 225ms post-DP: β = 7.39, *SE* = 1.02, lCI = 5.65, uCI = 9.72, BF01 < 0.01; 300ms post-DP: β = 8.04, *SE* = 1.88, lCI = 4.99, uCI = 12.32, BF01 < 0.01). Figure 3D shows that, with the gating segment becoming longer, the rating scores gradually reduce (lower scores indicating more accurate and more confident identification). We examined whether the mean score at the DP is equal to 3.5 (i.e. chance performance) and found strong evidence supporting the null hypothesis, β = −0.02, *SE* = 0.04, lCI = −0.10, uCI = 0.06, BF01 = 21.79, which is consistent with the accuracy results. Furthermore, in order to refine the estimate of the time point at which participants recognise the stimuli with enough confidence, we also investigated at what alignment point is there evidence showing the mean score lower than 2 (i.e. participants indicating more confident identification). We found moderate evidence supporting the null hypothesis (mean score equals to 2) at 75ms post-DP (β = −0.09, *SE* = 0.08, lCI = −0.25, uCI = 0.07, BF01 = 6.07), but extremely strong evidence in favour of the alternative hypothesis at 150ms post-DP (β = −0.71, *SE* = 0.05, lCI = −0.79, uCI = 0.62, BF01 < 0.01). These results show that critical acoustic information that supports confident word recognition arrives between 75ms and 150ms post-DP.

Overall, the post-test gating study confirmed that the pre-DP segments of correctly morphed stimuli are not distinguishable within each stimuli set. However, since we found items that were not correctly morphed during this control study, we did a thorough check of our stimuli and identified all the problematic items (16 words and 12 pseudowords), which resulted in 8.68% of all trials presented in the MEG study. In order to double check our MEG study results, we then removed all these problematic trials from the data and reanalysed the data using the same methods as described in the method section. Fortunately, we did not find any inconsistent pattern or significance in our behavioural or neural results compared to those reported with all trials included. Therefore, we kept the original MEG and behavioural results with all items included in this paper.

### Behavioural Data Analyses

Response times (RTs) were measured from the onset of the stimuli and inverse-transformed so as to maximise the normality of the data and residuals; Figures report untransformed response times for clarity. Inverse-transformed RTs and error rates were analysed using linear and logistic mixed-effect models respectively using the lme4 package in R (Bates et al. 2014). Lexicality (word, pseudoword) and prime type (unprimed, primed by same lexical status, primed by different lexical status) were fixed factors, while participant and item were random factors. Maximal models accounting for all random effects were attempted wherever possible, but reduced random effects structures were applied when the full model did not converge (Barr et al., 2013). Likelihood-ratio tests comparing the full model to a nested reduced model using the Chi-Square distribution were conducted to evaluate main effects and interactions. Significance of individual model coefficients were obtained using *t* (reported by linear mixed-effect models) or *z* (reported by logistic mixed-effect models) statistics in the model summary. One-tailed *t* statistics for RTs are also reported for two planned contrasts: (1) word-primed versus unprimed conditions for word targets, and (2) word-primed versus pseudoword-primed conditions for word targets.

When assessing priming effects, we excluded data from target trials in which the participant made an error in the corresponding prime trial, because it is unclear whether such target items will be affected by priming given that the prime word was not correctly identified. In addition, three trials with RTs shorter than the average pre-DP length (353ms) were removed from further analysis, since responses before words and pseudowords acoustically diverge are too quick to be valid lexical decision responses.

### MEG Data Acquisition, Processing and Analyses

Magnetic fields were recorded with a VectorView system (Elekta Neuromag) which contains a magnetometer and two orthogonal planar gradiometers at each of 102 locations within a hemispherical array around the head. Although electric potentials were recorded simultaneously using 68 Ag-AgCl electrodes according to the extended 10-10% system, these EEG data were excluded from further analysis due to excessive noise. All data were digitally sampled at 1 kHz. Head position were monitored continuously using five head-position indicator (HPI) coils attached to the scalp. Vertical and horizontal electro-oculograms were also recorded by bipolar electrodes. A 3D digitizer (FASTRAK; Polhemus, Inc.) was used to record the positions of three anatomical fiducial points (the nasion, left and right preauricular points), HPI coils and evenly distributed head points for use in source reconstruction.

MEG Data were preprocessed using the temporal extension of Signal Source Separation in MaxFilter software (Elekta Neuromag) to reduce noise sources, normalise the head position over blocks and participants to the sensor array and reconstruct data from bad MEG sensors. Subsequent processing was conducted in SPM12 (https://www.fil.ion.ucl.ac.uk/spm/) and FieldTrip (http://www.fieldtriptoolbox.org/) software implemented in MATLAB. The data were epoched from −1100 to 2000ms time-locked to the DP and baseline corrected relative to the −1100 to −700ms prior to the DP, which is a period before the onset of speech for all stimuli (Figure 1C). Low-pass filtering to 40 Hz was conducted both before and after robust averaging across trials (Litvak et al., 2011). A time window of −150 to 0ms was defined for pre-DP comparisons based on the shortest pre-DP stimuli length. A broad window of 0 to 1000ms was defined for post-DP comparisons, which covered the possible period for lexicality and prime effects. After averaging over trials, an extra step was taken to combine the gradiometer data from each planar sensor pair by taking the root-mean square (RMS) of the two amplitudes.

Sensor data from magnetometers and gradiometers were analysed separately. We converted the sensor data into 3D images (2D sensor x time) and performed *F* tests for main effects across sensors and time (the term “sensors” denotes interpolated sensor locations in 2D image space). Reported effects were obtained with a cluster-defining threshold of *p* < .001, and significant clusters identified as those whose extent (across space and time) survived *p* < 0.05 FWE-correction using Random Field Theory (Kilner & Friston, 2010). Region of interest (ROI) analyses for the priming effect were then conducted over sensors and time windows that encompassed the significant pseudoword>word cluster, orthogonal to priming effects. When plotting waveforms and topographies, data are shown for sensors nearest to the critical points in 2D image space.

Apart from the two planned contrasts mentioned above (see Behavioural Data Analyses), which were applied to post-DP analysis, one-tailed *t* statistics was also reported on the pre-DP planned contrast between unprimed and word-primed items.

### Source Reconstruction

In order to determine the underlying brain sources underlying the sensor-space effects, source reconstruction was conducted using SPM’s Parametric Empirical Bayes framework (Henson et al., 2011). To begin with, we obtained T1-weighted structural MRI (sMRI) scans from each participant on a 3T Prisma system (Siemens, Erlangen, Germany) using an MPRAGE sequence. The scan images were segmented and normalised to an MNI template brain in MNI space. The inverse of this spatial transformation was then used to warp canonical meshes derived from that template brain back to each subject’s MRI space (Mattout et al., 2007). Through this procedure, canonical cortical meshes containing 8196 vertices were generated for the scalp and skull surfaces. We coregistrated the MEG sensor data into the sMRI space for each participant by using their respective fiducials, sensor positions and head-shape points (with nose points removed due to the absence of the nose on the T1-weighted MRI). Using the single shell model, the lead field matrix for each sensor was computed for a dipole at each canonical cortical mesh vertex, oriented normal to the local curvature of the mesh.

Source inversion was performed with all conditions pooled together using the ‘IID’ solution, equivalent to classical minimum norm, fusing the magnetometer and gradiometer data (Henson et al, 2011). The resulting inversion was then projected onto wavelets spanning frequencies from 1 to 40 Hz and from −150 to 0ms time samples for pre-DP analysis and 400 to 900ms for post-DP analysis. This post-DP time window was defined by overlapping temporal extent of the pseudoword > word cluster between gradiometers and magnetometers. The total energy within these time-frequency windows was summarised by taking the sum of squared amplitudes, which was then written to 3D images in MNI space.

Reported effects for source analyses were obtained with a cluster-defining threshold of *p* < 0.05 (FWE-corrected). And as in sensor space, ROI analyses were conducted over significant sensors and time windows from the orthogonal pseudoword>word cluster. Factorial ANOVA were carried out on main effects and one-tailed paired *t*-tests on planned contrasts (see MEG Data Acquisition and Processing).

## Results

### Behaviour

#### Response Times

As shown in Figure 4A, factorial analysis of lexicality (word, pseudoword) and prime type (unprimed, primed by same lexical status, primed by different lexical status) indicated a significant main effect of lexicality, in which RTs for pseudowords were significantly longer than for words, *X^2^*(3) = 23.60, *p* < .001. In addition, there was a significant interaction between lexicality and prime type, *X^2^*(2) = 10.73, *p* = .005. This interaction was followed up by two separate one-way models for words and pseudowords, which showed a significant effect of prime type for words, *X^2^*(2) = 10.65, *p* = .005, but not for pseudowords, *X^2^*(2) = 1.62, *p* = .445. Consistent with the competitor priming results from Monsell and Hirsh (1998), words that were primed by another word sharing the same initial segments were recognised significantly more slowly than unprimed words (for mean raw RTs see Fig 3A), β = 0.02, *SE* = 0.01, *t*(79.69) = 3.33, *p* < .001, and more slowly than pseudoword-primed words, β = 0.02, *SE* = 0.01, *t*(729.89) = 2.37, *p* = .018. As mentioned earlier (see Introduction), both competitive- and predictive-selection models predicted longer response times to word-primed target words compared to unprimed words, it is hence critical to distinguish the two accounts through further investigation of the MEG responses.

**Figure 4.**
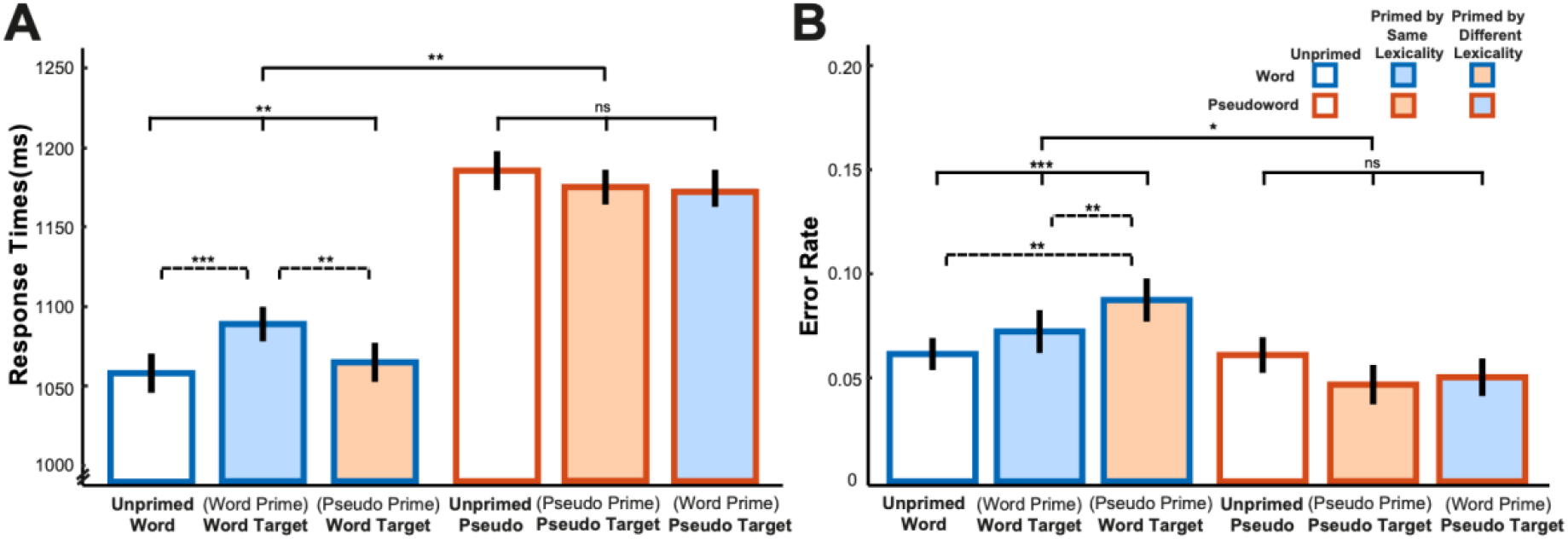
Response time results (***A***) and accuracy results (***B***) of the lexical decision task. Bars are color-coded by lexicality and prime type on the *x* axis (words, blue frame; pseudowords, orange frame; unprimed, no fill; primed by same lexicality, consistent fill and frame colors; primed by different lexicality, inconsistent fill and frame colors). Bars show the subject grand averages, error bars represent ± within-subject SE, adjusted to remove between-subjects variance (Cousineau, 2005). Statistical significance is shown based on generalised linear mixed-effects regression: * *p*<0.05, ** *p*<0.01, *** *p*<0.001. Statistical comparisons shown with solid lines indicate the lexicality by prime-type interaction and main effects of prime-type for each lexicality, whereas comparisons with broken lines indicate the significance of pairwise comparisons.

#### Accuracy

Figure 4B shows that there was a trend towards more lexical decision errors in response to words than to pseudowords, although this lexicality effect was marginal, *X^2^*(3) = 7.31, *p* = .063. The error rates for words and pseudowords were also affected differently by priming, as indicated by a significant interaction between lexicality and prime type, *X^2^*(2) = 6.08, *p* = .048. Follow-up analyses using two separate models for each lexicality type showed there was a main effect of prime type for words, *X^2^*(2) = 13.95, *p* < .001, but not for pseudowords, *X^2^*(2) = 1.93, *p* = .381. Since we had not anticipated these priming effects on accuracy, post-hoc pairwise *z* tests were Bonferroni corrected for multiple comparisons. These showed that pseudoword priming reliably increased the error rate compared to the unprimed condition, β = 1.68, *SE* = 0.54, *z* = 3.14, *p* = .005, and to the word-primed condition, β = 2.74, *SE* = 0.89, *z* = 3.07, *p* = .007. Although no specific predictions on accuracy were made a priori by either competitive- or predictive-selection model, it is worth noting that participants might have expected pseudowords to be repeated given the increased error rate of responses to pseudoword-primed target words.

### MEG

In order to explore the impact of lexicality and competitor priming on neural responses to critical portions of speech stimuli, both before and after they diverge from each other, MEG responses were time-locked to the DP. All reported effects are family-wise error (FWE)-corrected at cluster level for multiple comparisons across scalp locations and time at a threshold of *p* < 0.05. We reported data from gradiometers, magnetometers and source space wherever possible, since sensor x time analyses help define the time-windows used by source localisation. Although some minor effects were shown in only one of these analyses, our most interesting effects are reliable in all three data types.

#### Pre-DP analyses

We assessed neural responses before the DP, during which only the shared speech segments have been heard and hence the words and pseudowords in each stimulus set are indistinguishable. Since there could not have been any effect of lexical status pre-DP, only prime type effects were considered in this analysis. Predictive- and competitive-selection accounts make opposite predictions for pre-DP neural signals evoked by word-primed items compared to unprimed items. We therefore conducted an F-test for neural differences between these two conditions across the scalp and source spaces over a time period of −150 to 0ms before the DP. A significant cluster of 295 sensor x time points (*p* = .023) was found in gradiometers over the mid-left scalp locations from −28 to −4ms (Figure 5A), in which unprimed items evoked significantly greater neural responses than word-primed items. On the suggestion of a reviewer, and mindful of the potential for these pre-DP neural responses to be modulated by post-DP information, we report an additional analysis with a lengthened analysis time window of −150ms to 100ms. Again, we found a significant unprimed > word-primed cluster of 313 sensor x time points (*p* = .033) over the exact same locations in gradiometers from −28 to −3ms pre-DP, which confirmed that this pre-DP effect was not pushed forward by any post-DP effect. We did not find any cluster showing stronger neural responses for word-primed items than unprimed items and no clusters survived correction for multiple comparisons for magnetometer responses or for analysis in source space.

**Figure 5.**
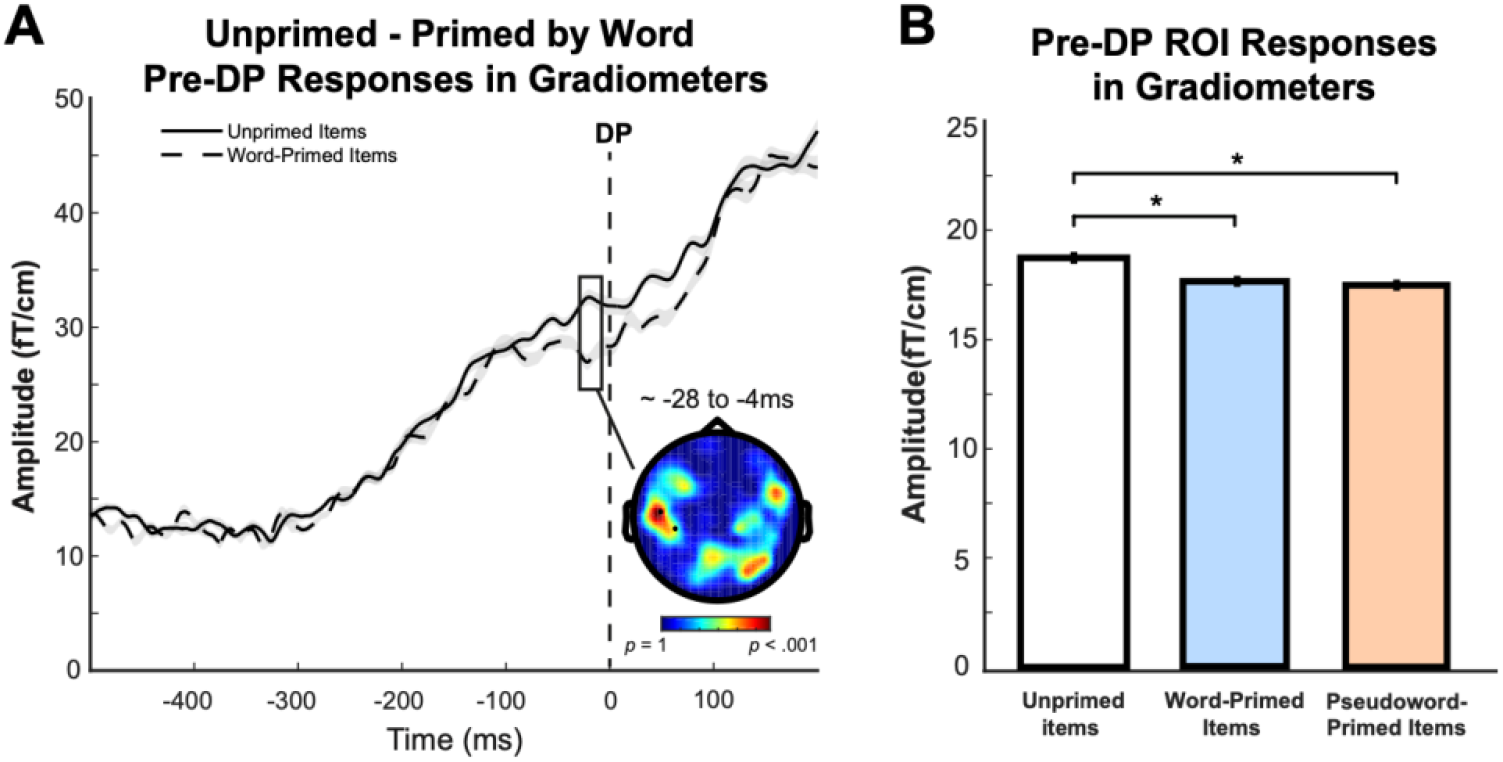
Pre-DP results. ***A* & *B.*** Pre-DP response difference between items that are unprimed and primed by a word in MEG gradiometer sensors within −150 to 0ms (a time window at which words and pseudowords are indistinguishable). The topographic plots show F-statistics for the entire sensor array with the scalp locations that form two statistically significant clusters highlighted and marked with black dots. Waveforms represent MEG response averaged over the spatial extent of the significant cluster shown in the topography. The grey shade of waveforms represents ± within-participant SE, adjusted to remove between-participants variance (Cousineau, 2005). ***C.*** ROI analysis of neural responses evoked by unprimed and primed items averaged over the same pre-DP time period of −150-0ms but across gradiometer sensor locations which showed the post-DP pseudoword>word lexicality effect (see Figure 5A). Bars are color-coded by prime type on the *x* axis (unprimed items, no fill; word-primed items, blue; pseudoword-primed items, orange; black frame indicates that words and pseudowords are indistinguishable). All error bars represent ± within-participant SE, adjusted to remove between-participant variance. Statistical significance: * *p*<0.05.

To further examine these results, we also conducted ROI analysis of gradiometer signals evoked by unprimed and primed items averaged over the same −150 to 0ms pre-DP time window but across the scalp locations that showed the post-DP lexicality effect at which pseudowords elicited greater neural responses than words (see Figure 6A). As shown in Figure 5B, the results indicated that unprimed items elicited significantly stronger neural responses than word-primed items, *t*(21) =2.41, *p* = .013, consistent with the whole-brain analysis. In particular, the mid-left cluster shown in panel A partially overlaps with the post-DP pseudoword>word cluster. The direction and location of these pre-DP neural responses are in accordance with the predictive-selection account and inconsistent with the competitive-selection account. A surprising finding is that post-hoc analysis also showed greater neural responses evoked by unprimed items than pseudoword-primed items, *t*(21) = 2.69, *p* = .014, although we had not predicted these effects from pseudoword primes.

**Figure 6.**
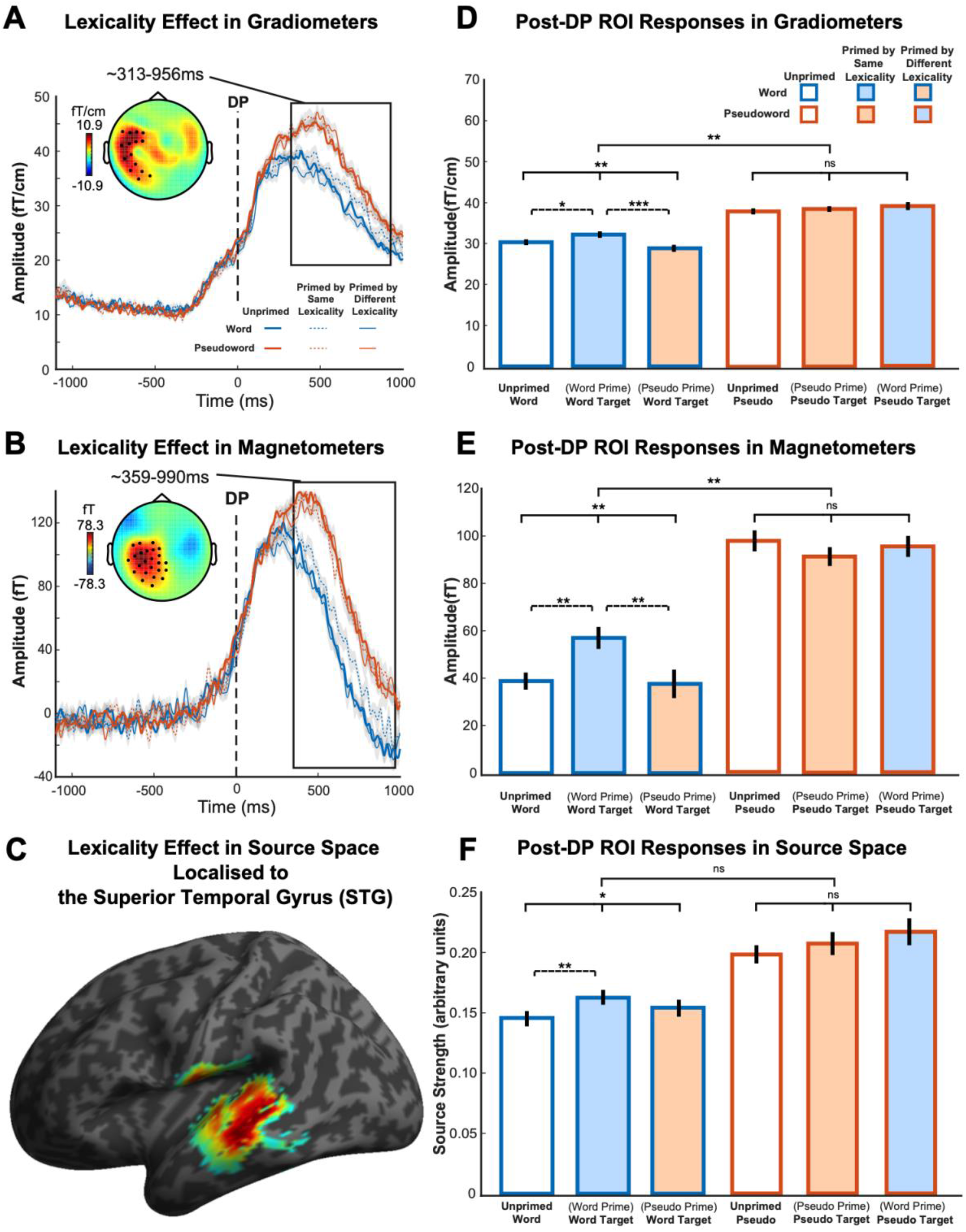
Post-DP results showing lexicality effects and corresponding ROI responses evoked by conditions of interest. ***A* & *B*.** Post-DP lexicality effects in MEG gradiometer and magnetometer sensors. The topographic plots show the statistically significant cluster with a main effect of lexicality (pseudoword > word). Waveforms represent MEG response averaged over the spatial extent of the significant cluster shown in the topography. The grey shade of waveforms represents ± within-participant SE, adjusted to remove between-participants variance. ***C.*** Statistical parametric map showing the cluster (pseudoword > word) rendered onto an inflated cortical surface of the Montreal Neurological Institute (MNI) standard brain thresholded at FWE-corrected cluster-level *p* < 0.05, localised to the left STG. ***D, E & F.*** Post-DP ROI ANOVA on neural signals and source strength evoked by conditions of interest averaged over the time window and scalp locations of the significant cluster shown in panel A, B & C. Bars are color-coded by lexicality and prime type on the *x* axis (words, blue frame; pseudowords, orange frame; unprimed, no fill; primed by same lexicality, consistent fill and frame colors; primed by different lexicality, inconsistent fill and frame colors). All error bars represent ± within-participant SE, adjusted to remove between-participants variance. Statistical significance from ANOVAs: * *p*<0.05, ** *p*<0.01, *** *p*<0.001. Statistical comparisons shown with solid lines indicate the lexicality by prime-type interaction and main effects of prime-type for each lexicality, whereas comparisons with broken lines indicate the significance of planned pairwise comparisons.

#### Post-DP analyses

We then examined the post-DP response differences between words and pseudowords (lexicality effect). The gradiometer sensors showed a significant cluster of 39335 sensor x time points (*p* < .001) over the left side of the scalp at 313-956ms post-DP (Figure 6A). In this cluster, pseudowords evoked a significantly stronger neural response than words. Similarly, magnetometer sensors also detected a significant left-hemisphere cluster of 68517 sensor x time points (*p* < .001) at 359-990ms post-DP (Figure 6B) showing the same lexicality effect. We did not find any significant cluster in which words evoked greater neural responses than pseudowords. These results are consistent with findings from Gagnepain and colleagues (2012). To locate the likely neural source of the effects found in sensor space, we conducted source reconstruction by integrating gradiometers and magnetometers. As shown in Figure 6C, results from source space showed that neural generators of the lexicality effect were estimated to lie within the left superior temporal gyrus (STG, volume of 2315 voxels, *p* < .001, peak at x = −46, y = −36, z = 0; x = −52, y = −34, z = −6; x = −56, y = −28, z = −10). This location, and direction of response, is consistent with a sub-lexical (e.g. phonemic) process being modulated by lexicality; in line with the predictive-selection account.

Next, we investigated whether the neural responses that were modulated by lexicality were also influenced by prime type by conducting an ROI analysis which tested the interaction between prime type and lexicality, as well as planned pairwise comparisons of priming effects on words alone, using data averaged over the time window and the sensor locations of the significant cluster shown in panel A and B (Figure 6D & E). Since these planned pairwise comparisons involve responses to familiar words only (i.e. words that are word-primed vs unprimed, words that are word-primed vs pseudoword-primed), they are orthogonal to the lexicality effect that defined the pseudoword>word cluster and hence are not confounded by task. The interaction was significant in both gradiometers, *F*(1.96, 41.11) = 7.30, *p* = .002, and magnetometers, *F*(1.90, 39.99) = 5.80, *p* = .007. Specifically, there was a significant effect of prime type for words, *F*(1.93, 40.55) = 8.01, *p* = .001 (gradiometers), *F*(1.81, 37.96) = 5.61, *p* = .009 (magnetometers), such that neural signals evoked by word-primed words were significantly stronger than those evoked by unprimed words, *t*(21) = 2.22, *p* = .019 (gradiometers), *t*(21) = 3.33, *p* = .002 (magnetometers), and pseudoword-primed words, *t*(21) = 3.70, *p* < .001 (gradiometers), *t*(21) = 2.64, *p* = .008 (magnetometers). In contrast, there was no reliable main effect of prime type for pseudowords, *F*(1.94, 40.80) = 0.67, *p* = .514 (gradiometers), *F*(1.79, 37.61) = 0.80, *p* = .446 (magnetometers). The corresponding tests performed on the source-reconstructed power within the lexicality ROI of suprathreshold voxels (Figure 6F) did not show a reliable interaction effect between lexicality and competitor priming, *F*(1.56, 32.85) = 0.99, *p* = .36. Nevertheless, consistent with sensor space results, source power did show a significant effect of prime type for words, *F*(1.73, 36.42) = 3.77, *p* = .038, but not pseudowords, *F*(1.62, 33.94) = 1.12, *p* = .326. Pairwise comparisons also indicated that word-primed words evoked significantly greater source strength than unprimed words, *t*(21) = 2.66, *p* = .007, though the effect between word-primed and pseudoword-primed words was not significant, *t*(21) = 1.26, *p* = .110. Overall, in line with behavioural results, neural responses evoked by words and pseudowords were also influenced differently by prime type. Critically, competitor priming modulated the post-DP neural responses evoked by words, but not those evoked by pseudowords, and these effects were localised to the left STG regions that plausibly contribute to sub-lexical processing of speech. This matches the pattern of responses proposed in the predictive-selection model (see Figure 1F).

As encouraged by a reviewer, we also conducted whole brain analyses for the competitor priming effects. We found a significant word-primed word > unprimed word cluster of 1197 sensor x time points (*p* = .034) in magnetometers in the left hemisphere within a time window of 426 - 466ms post-DP. We also found a significant and a marginal word-primed word > pseudoword-primed word cluster in gradiometers in the left hemisphere respectively of 527 sensor x time points (*p* = .011) at 719-749ms and 471 sensor x time points (*p* = .053) at 315-336ms. These topographies and time courses overlap with the pseudoword > word clusters and are consistent with our ROI results. Hence, the ROI analyses have picked up the most important findings from these whole-brain analyses.

To ensure that other response patterns were not overlooked, we also investigated whether there was any lexicality by prime-type interaction at other locations across the scalp and source spaces, and during other time periods. As shown in Figure 7A, a significant cluster of Gradiometers at midline posterior scalp locations were found at 397-437ms post-DP, in which the effect of priming was significantly different for words and pseudowords. Figure 7B shows gradiometer signals evoked by conditions of interest averaged over the spatial and temporal extent of the significant cluster in panel A. To explore this profile, we computed an orthogonal contrast to assess the overall lexicality effect (the difference between words and pseudowords), and the result was marginal, *F*(1.00, 21.00) = 3.50, *p* = .075. The effect of prime type was marginally significant for words, *F*(1.89, 39.78) = 3.08, *p* = .060, but significant for pseudowords, *F*(1.80, 37.85) = 7.14, *p* = .003. The location and pattern of this interaction cluster were dissimilar to those predicted by either competitive- or predictive-selection theories and no cluster survived correction in magnetometer sensors or source space hence we did not consider this effect to be as relevant or interpretable as our other findings. We report it here in the interest of completeness and transparency.

**Figure 7.**
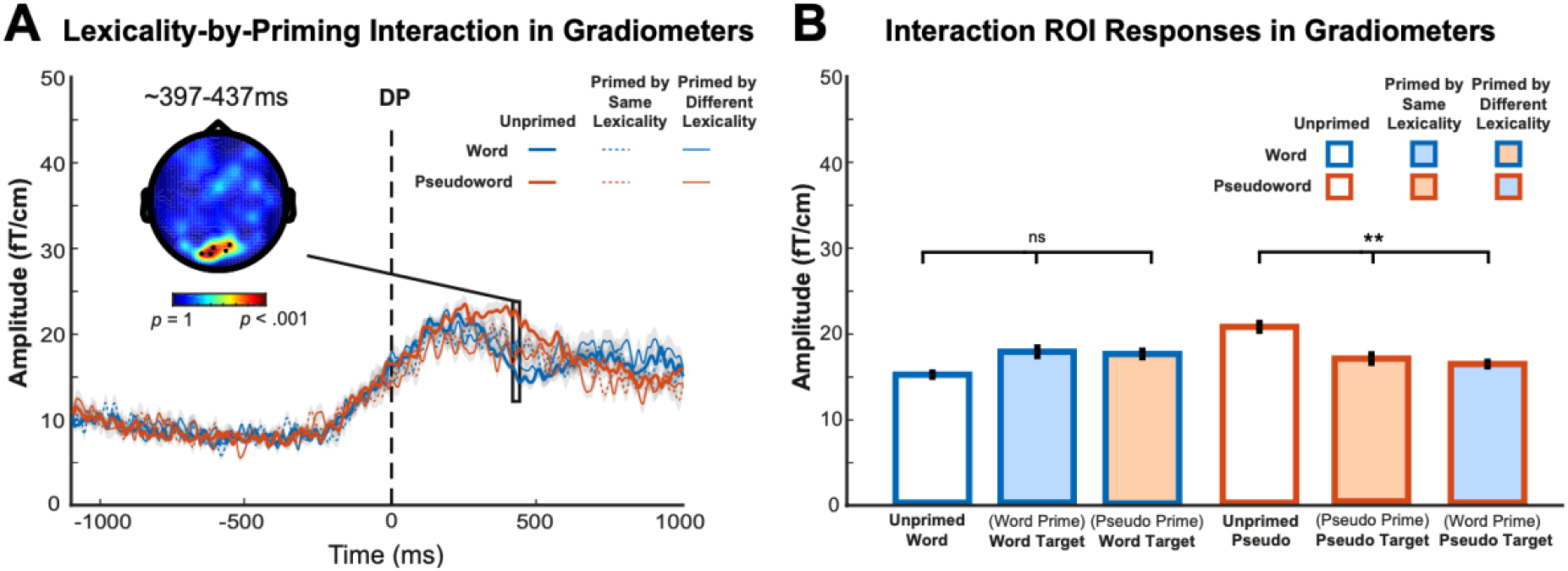
Post-DP results showing lexicality-by-priming interaction effects in MEG gradiometers. ***A.*** The topographic plot shows *F*-statistics for the statistically significant cluster that showed an interaction between lexicality and prime type. Waveforms represent gradiometer responses averaged over the spatial extent of the significant cluster shown in the topography. The grey shade of waveforms represents ± within-participant SE, adjusted to remove between-participants variance. ***B***. Gradiometer signals evoked by conditions of interest averaged over temporal and spatial extent of the significant cluster in panel A. All error bars represent ± within-participant SE, adjusted to remove between-participants variance. Statistical significance: ** *p*<0.01. The statistical comparison lines indicate main effects of prime type for each lexicality. The lexicality by prime-type interaction is statistically reliable as expected based on the defined cluster.

#### Linking neural and behavioural effects

To further examine the relationship between neural and behavioural response differences attributable to competitor priming or lexicality, we conducted a single-trial regression analyses using linear mixed-effect models that account for random intercepts and slopes for participants and stimuli sets (grouped by their initial segments). We calculated behavioural RT differences and neural MEG differences caused by: (1) lexicality. i.e. the difference between pseudoword and word trials (collapsed over primed and unprimed conditions) and (2) competitor priming, i.e. the difference between unprimed and word-primed word trials, with MEG signals averaged over the spatial and temporal extent of the post-DP pseudoword>word cluster seen in sensor space and the STG peak voxel in source space (see Figure 6). We then assessed the relationship between these behavioural and neural difference effects in linear mixed-effect regression of single trials, with differences in RTs as the independent variable and differences in MEG responses as the dependent variable. The analyses were conducted using the lme4 package in R (Bates et al. 2014).

As shown in Figure 8A, we observed a significant positive relationship between RTs and magnetometers on lexicality difference (β = 0.11, *SE* = 0.01, *t*(23.31) = 7.77, *p* < .001), although associations between RTs and gradiometers or source response were not significant. These observations from magnetometers indicated that slower lexical decision times evoked by pseudowords were associated with greater neural responses. Furthermore, the intercept parameter for the magnetometers model was significantly larger than zero, β = 37.58, *SE* = 5.72, *t*(23.09) = 6.57, *p* < .001. We can interpret this intercept as the neural difference that would be predicted for trials in which there was no delayed response to pseudowords compared to words. The significant intercept indicated a baseline difference in neural responses to words and pseudowords, even in the absence of any difference in processing effort (as indexed by lexical decision RTs). This suggested the engagement of additional neural processes specific to pseudowords regardless of the behavioural effect (cf. Taylor et al., 2014).

**Figure 8.**
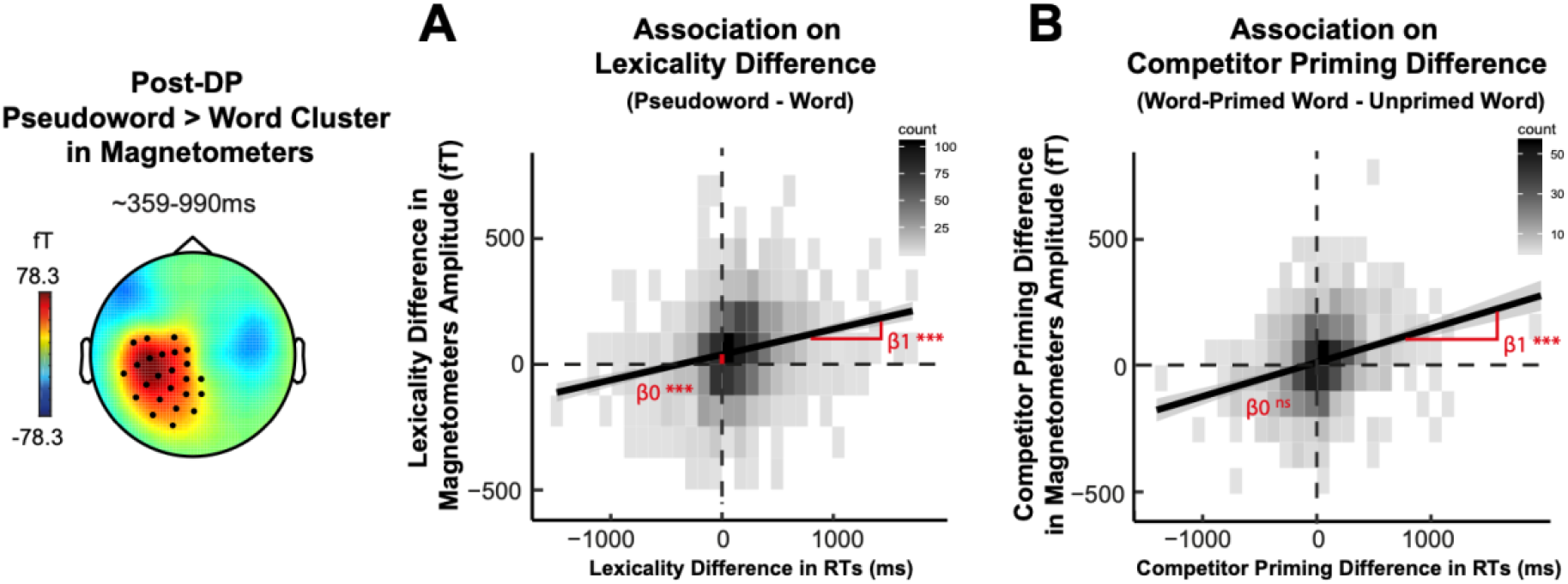
Single-trial linear mixed-effect models which accounted for random intercepts and slopes for participants and stimuli sets (grouped by initial segments) were constructed to compute the relationship between RTs and magnetometers on **(*A*)** lexicality difference (i.e. between pseudowords and words, collapsed over unprimed and primed conditions) and **(*B*)** competitor priming difference (i.e. between word-primed word and unprimed word conditions). Magnetometer responses were averaged over the time window and scalp locations of the significant post-DP pseudoword>word cluster (see Figure 6). β1 refers to the model slope, β0 refers to the model intercept. Statistical significance: *** *p*<0.001.

Figure 8B showed another significant positive relationship between RTs and magnetometers on competitor priming difference (β = 0.15, *SE* = 0.02, *t*(38.85) = 7.89, *p* < .001), while relationships between RTs and gradiometers or source response were again not significant. Interestingly, unlike for the lexicality effect, the intercept in this competitor priming magnetometers model did not reach significance (β = 12.88, *SE* = 7.27, *t*(21.33) = 1.77, *p* = .091). This non-significant intercept might suggest that if word-primed words did not evoke longer RTs than unprimed words, magnetometer signals would not be reliably different between the two conditions either. Hence, consistent with predictive-selection accounts, the increased post-DP neural responses in the STG caused by competitor priming was both positively linked to and mediated by longer response times.

## Discussion

In this study, we distinguished different implementations of Bayesian perceptual inference by manipulating the prior probability of spoken words and examining the pattern of neural responses. We replicated the competitor priming effect such that a single prior presentation of a competitor word (e.g. *hijack*) delayed the recognition of a similar-sounding word (e.g. *hygiene*), whereas this effect was not observed when the prime or target was a pseudoword (e.g. *hijure*). Armed with this behavioural evidence, we used MEG data to test the neural bases of two Bayesian theories of spoken word recognition.

### Competitive-vs predictive-selection

Competitive-selection accounts propose that word recognition is achieved through direct inhibitory connections between representations of similar candidates (e.g. McClelland & Elman, 1986). Priming boosts the activation of heard words and increases lateral inhibition applied to neighbouring words, which delays their subsequent identification. The effect of competitor priming is to increase lexical uncertainty, and hence lexical-level neural responses, until later time points when target words can be distinguished from the competitor prime (Figure 1C). In contrast, predictive-selection accounts propose that word recognition is achieved by subtracting predicted speech from heard speech and using computations of prediction error to update lexical probabilities (Davis & Sohoglu, 2020). By this view, predictions for segments that are shared between competitor primes and targets (pre-DP segments) will be enhanced after presentation of prime words. Thus, competitor priming will reduce the magnitude of prediction error, and hence neural responses pre-DP (Figure 1F). Only when speech diverges from predictions (post-DP segments) will competitor-primed words evoke greater prediction error, leading to increased neural response in brain areas involved in pre-lexical (e.g. phonemic) processing of speech representing prediction error (Blank et al., 2018; Blank & Davis, 2016).

It should be acknowledged that both models involve multiple levels of representation and hence both sub-lexical and lexical processes. However, our focus is on lexical processing within the competitive-selection framework and sub-lexical processing within the predictive-selection framework. These are the critical levels that 1) support word recognition according to each theory, 2) are modulated by the competitor priming effect that our study manipulates and 3) are invoked to explain the slower behavioural responses and associated changes in MEG responses that we observed.

We tested the predictions for the direction and timing of neural responses associated with competitor priming using MEG data which showed opposite neural effects pre- and post-DP. In the pre-DP period, consistent with predictive-selection but contrary to competitive-selection mechanism, we saw decreased neural responses for word-primed items compared to unprimed items. The initial, shared segments between prime (*hijack*) and target (*hygiene*) words evoked a reduced response during early time periods in line with a reduction in prediction error. However, during the post-DP period, we found competitor-primed words evoked stronger neural responses than unprimed words in exactly the same locations and time periods that showed increased responses to pseudowords (*hijure*) compared to words. These post-DP response increases are in line with enhanced processing difficulty for competitor-primed words and pseudowords due to greater prediction error. Thus, the time course of the competitor priming neural effects – showing reduced neural responses pre-DP and increased neural responses post-DP – closely resembles the expected changes in prediction error (Figure 1F) based on predictive-selection mechanisms.

On top of the direction and timing of neural responses, effects of lexicality and competitor priming localised to the left STG. This is a brain region that has long been associated with lower-level sensory processing of speech (Yi et al., 2019). Our observation of increased responses to pseudowords in this region is in accordance with source-localised MEG findings (Gagnepain et al., 2012; Shtyrov et al., 2012) and evidence from a meta-analysis of PET and fMRI studies (Davis & Gaskell, 2009). This location is also consistent with the proposal that lexical influences on segment-level computations produce reliable neural differences between words and pseudowords (Davis & Sohoglu, 2020). We take this finding as further evidence in favour of computations of segment prediction error as a critical mechanism underlying word identification.

We further show using regression analyses that neural (MEG) and behavioural (RT) effects of lexicality and competitor priming are linked on a trial-by-trial basis. Trials in which pseudoword processing or competitor priming leads to larger increases in RT also have greater post-DP neural responses. These links between behavioural and neural effects of lexicality and competitor priming are once more in-line with the proposal that post-DP increases in prediction error are a key neural mechanism for word and pseudoword processing and can explain the delayed behavioural responses seen in competitor priming. Interestingly, lexicality and competitor priming effects differ in terms of whether a reliable neural response difference would be seen for trials with no baseline RT difference. While neural lexicality effects were significant even for trials that did not show behavioural effects, the same was not true for the competitor priming effect. These results indicate that, consistent with predictive-selection accounts, the post-DP neural competitor priming effect was mediated by changes in behavioural RTs. In contrast, an increased neural response to pseudowords was expected even in trials for which RTs did not differ between pseudowords and words. We will consider the implications of these and other findings for pseudoword processing in the next section.

### How do listeners process pseudowords?

Participants identified pseudowords with a speed and accuracy similar to that seen during recognition of familiar words. This is consistent with an optimally-efficient language processing system (Marslen-Wilson, 1984; Zhuang et al, 2014), in which pseudowords can be distinguished from real words as soon as deviating speech segments are heard. Beyond this well-established behavioural finding, however, we reported two seemingly contradictory observations concerning pseudoword processing.

The first is that, while post-DP neural activity and response times for words were modulated by competitor priming, processing of pseudowords was not similarly affected. This might suggest that the prior probability of hearing a pseudoword and the prediction error elicited by mismatching segments are not changed by our experimental manipulations. This may be because pseudowords have a low or zero prior probability and elicit maximal prediction errors that cannot be modified by a single prime. Yet, memory studies suggest that even a single presentation of a pseudoword can be sufficient for listeners to establish a lasting memory trace (Mckone & Trynes, 1999; Arndt et al., 2008). However, it is possible that this memory for pseudowords reflects a different type of memory (e.g. episodic memory) from that produced by a word, with only the latter able to temporarily modify long-term, lexical-level representations and predictions for word speech segments (cf. Complementary Learning Systems theories, McClelland et al., 1995; Davis & Gaskell, 2009).

A second observation is that, contrary to the null result for post-DP processing, pseudoword priming reduced subsequent pre-DP neural responses evoked by target items to a similar degree as word priming (Figure 5B). This pre-DP effect is surprising given previous evidence suggesting that pseudowords must be encoded into memory and subject to overnight, sleep-associated consolidation in order to modulate the speed of lexical processing (Tamminen et al., 2010; James et al., 2017) or neural responses (Davis & Gaskell, 2009; Landi et al. 2018). It might be that neural effects seen for these pre-DP segments were due to changes to the representation of familiar words that our pseudowords resembled, though these were insufficient to modulate processing of post-DP segments.

### Summary

Our work provides compelling evidence in favour of neural computations of prediction error during spoken word recognition. Although the previous work by Gagnepain et al. (2012) also provided evidence for the predictive-selection account, their behavioural effects of consolidation on word recognition were obtained during different tasks and different sessions from their neural responses. Our current study goes beyond this previous work by adopting a single task (lexical decision) and using a competitor priming paradigm that permits concurrent measurement of perceptual outcomes and neural responses in a single session. This enables us to directly link trials that evoked stronger neural signals in the STG to delayed RTs and hence provide stronger evidence that both of these effects are caused by competitor priming.

In addition, unlike previous work (Brodbeck et al. 2018; Donhauser & Baillet, 2020) which reported neural responses correlated with lexical entropy as well as prediction error (surprisal), we did not find any similarly equivocal evidence. These earlier studies measured neural responses to familiar words in continuous speech sequences such as stories or talks. It might be that effects of lexical entropy are more apparent for connected speech than isolated words. However, since lexical uncertainty (entropy) and segment-level predictability (segment prediction error or surprisal) are highly correlated in natural continuous speech, these studies may be less able to distinguish between the lexical and segmental mechanisms that we assessed here. In contrast, our speech materials were carefully selected to change lexical probability (through priming) and for priming to have opposite effects on segment prediction error before and after DP. This manipulation provides evidence in favour of predictive-selection mechanisms that operate using computations of prediction error during spoken word recognition.

## Acknowledgments

The research was supported by the UK Medical Research Council (SUAG/044 & SUAG/046 G101400) and by a China Scholarship Council award to Yingcan Carol Wang. We are grateful to Clare Cook, Ece Kocagoncu and Tina Emery for their assistance with data collection, and also to Olaf Hauk for his advice on MEG data analysis.

